# Algorithmic Assessment of Missense Mutation Severity in the Von-Hippel Lindau Protein

**DOI:** 10.1101/2020.05.20.106021

**Authors:** Francisco R. Fields, Niraja Suresh, Morgan Hiller, Stefan D. Freed, Kasturi Haldar, Shaun W. Lee

## Abstract

Von Hippel-Lindau disease (VHL) is an autosomal dominant rare disease that causes the formation of angiogenic tumors. When functional, pVHL acts as an E3 ubiquitin ligase that negatively regulates hypoxia inducible factor (HIF). Genetic mutations that perturb the structure of pVHL result in dysregulation of HIF, causing a wide array of tumor pathologies including retinal angioma, pheochromocytoma, central nervous system hemangioblastoma, and clear cell renal carcinoma. These VHL-related cancers occur throughout the lifetime of the patient, requiring frequent intervention procedures, such as surgery, to remove the tumors. Although VHL is classified as a rare disease (1 in 39,000 to 1 in 91,000 affected) there is a large heterogeneity in genetic mutations listed for observed pathologies. Understanding how these specific mutations correlate with the myriad of observed pathologies for VHL could provide clinicians insight into the potential severity and onset of disease. Using a set of 285 ClinVar mutations in VHL, we developed a multiparametric scoring algorithm to evaluate the overall clinical severity of missense mutations in pVHL. The mutations were assessed according to eight weighted parameters as a comprehensive evaluation of protein misfolding and malfunction. Higher mutation scores were strongly associated with pathogenicity. Our approach establishes a novel *in silico* method by which VHL-specific mutations can be assessed for their severity and effect on the biophysical functions of the VHL protein.

## Introduction

Von Hippel-Lindau (VHL) disease is an autosomal-dominant hereditary disease associated with the development of multiple angiogenic tumor types. This includes clear cell renal carcinoma (ccRCC), retinal angioma (RA), central nervous system hemangioblastoma (CHB), and pheochromocytoma (PCC)(1,2). The presence or absence of PCC divides VHL disease into type 1 or type 2. Type 2 VHL is further subdivided into three subtypes depending on the appearance of other cancers: type 2A, PCCs but no ccRCCs, type 2B, PCCs and ccRCCs, or type 2C, PCCs only(1). While this allows for some preliminary genotype-phenotype associations, a patient’s association with a specific subtype alternates as different cancers arise throughout their lifetime(1).

Patients with VHL disease have a single mutation in one allele of the *VHL* gene(3). Upon spontaneous inactivation of the second allele, tumor development can initiate, making the loss of heterozygosity (LOH) a crucial step in the development of VHL disease(1,4,5). The *VHL* gene encodes two protein products, both of which exhibit equivalent activity: the 30kDa isoform (pVHL_30_) and the more common 19kDa isoform (pVHL_19_) found in most tissues (6,7). pVHL forms a complex with elongin B (EloB) and elongin C (EloC) for the VCB complex(8–10). This stabilizes EloB, EloC, and pVHL, making them resistant to proteosomal degradation; however, upon mutation of pVHL, contacts with EloB and C become disrupted, making pVHL unstable and a target for degradation(9,11,12). VCB then complexes with cullin 2 (Cul2) and the RING finger protein RBX1 to form the VCB-CR complex(9). This complex functions as an E3 ubiquitin ligase, targeting a variety of proteins for degradation by the proteasome(13–15).

HIFα is ubiquitinated by the VCB-CR complex for degradation by the proteasome(2,13,16). HIF is involved in cellular oxygen sensing and regulates the expression of angiogenic genes making it a key player in the development of the vascularized tumor pathologies associated with VHL disease(16,17). Under normoxic conditons, HIF-1α is hydroxylated on two proline residues allowing for interaction with pVHL and its subsequent ubiquitination by the VCB-CR complex(2,16). In hypoxic conditions, HIF-1α is not hydroxylated, preventing negative regulation by pVHL. Under these conditions, active HIF1α subsequently drives the expression of hypoxia associated genes. Loss of functional pVHL allows aberrant expression of HIF target genes, such as vascular endothelial growth factor, contributing to the development of VHL associated angiogenic tumors (18–20).

Regardless of VHL subtype, patients are at a lifetime risk for the development of tumors with the age of onset of VHL disease ranging from 20 to 40 years old(21). Clinical diagnosis of VHL disease is dependent upon the familial history of VHL. Patients with a family history of VHL must present with CHB, PCC, or ccRCC; however, if there is no family history of disease, patients must then present with two more CHBs or a CHB and a visceral tumour, such as ccRCC(1,2,21). Genetic testing is conducted for presymptomatic detection of VHL for patients with a family history of disease(22). Surveillance, which varies since there are many tissue types in which the VHL tumors and cysts can arise, includes ophthalmologic evaluation and CT or MRI scans(21,23). Similar to surveillance, treatment is also varied due to the breadth of tumor types and includes surgery, radiation, or chemotherapies(21,23).

Multiple studies have investigated the association of mutation types to the VHL subtypes; however, there is still heterogeneity associated with the phenotypes of missense mutations(24–26). While loss-of-function mutations cause global disruption of the VHL protein, missense mutations may only affect certain interaction partners and cellular pathways involving pVHL(27). A recent study completed by Razafinjatovo et al used an *in silico* approach to determine the thermodynamic stability of a given pVHL mutation(28). It was determined that the most thermodynamically unstable missense mutations resulted in pathogenic disease via global destabilization of pVHL and stabilization of HIF. This suggests that while some VHL missense mutations might only affect specific functions of the protein, others cause global misfolding and destabilization of the protein. A comprehensive examination of the effects of a given missense mutation for pVHL can provide significant insight into how a given patient mutation can be predictive of disease severity; however, a systematic examination of the role of a given missense mutation (and subsequent amino acid replacement) must take into account multiple factors: secondary structure, thermodynamic stability, binding partners, translation rate, among other biophysical and biochemical properties. Providing a predictive scale of the phenotypic severity of a given missense mutation using *in silico* evaluation can potentially inform clinicians to develop tailored screening and surveillance strategies for each patient. Currently, some online databases provide investigators with basic information on the pathogenicity of a given mutation in genetic diseases, including VHL. *ClinVar* provides basic annotation on the pathogenicity of curated mutations according to the American College of Medical Genetics and Genomics (ACMG)(29,30). These guidelines provide a spectrum of pathogenicity descriptors for mendelian genetic diseases. Within these guidelines, mutations annotated as “pathogenic” or “likely pathogenic” have a greater than 90% certainty of a given gene variant being disease causing(30). Leveraging these sources of phenotypic information can help train and refine predictive algorithms for the assessment of missense mutation severity. Previously, we developed a computational, multiparameteric approach to evaluate the biophysical consequences of missense mutations on the structure and stability of the Mucopolysaccharidosis Type IIIA (Sanfilippo Syndrome) protein (MPSIIIA). Severe mutations identified through our scoring approach correlated to a higher clinical severity of Sanfilippo Syndrome(31). We observed that mutations more deleterious to overall enzyme folding and function were correlated to more severe disease outcomes and a higher multiparameteric algorithm scores(31). In this study we created an advanced weighted-score multiparametric approach to validate the use of a computational algorithm to assess the potential disease severity of genetic missense mutations in pVHL. We focused not only on mutations that can affect the overall proteostasis of pVHL, but also noted the specific mutations that would impact VHL-specific functional properties (27,28,32). Our multiparametric algorithm for VHL included a set of eight biophysical parameters with individually weighted scores that gave an overall assessment of the ability of a given missense mutation in VHL to result in protein impairment: 1. aggregation propensity; 2. protein-protein interactions; 3. secondary structure; 4. conformational flexibility; 5. solvent accessibility; 6. protein stability; 7. post-translational modifications, and 8. translational rate(9,14,17,31).

## Materials and Methods

### Mutation sets

A set of 285 missense mutations in the human VHL gene, arising from a single nucleotide polymorphism (SNP) was acquired from ClinVar(29) (Supplemental File 2). An additional set of 1380 mutations was generated to represent all possible theoretical missense mutations (APMM) of VHL from a SNP (Supplemental File 1). Finally, hot spot mutations and mutation lists associated with different pathogenic outcomes were selected from the literature(25,28,32–34) (Supplemental File 3). A total of 1665 mutations were therefore used in our multiparametric analysis.

### Structures used in Analysis

The VHL crystal structure in complex with EloB, EloC, and Cul2 was used in Parameters 2, 3, 5, and 6 (1VCB)(35). Crystal structures in complex with HIF-1a were also used to develop Parameter 2 (1LM8, 4WQO)(9,36). The unstructured N-terminus of VHL is missing from published crystal structures; therefore, to assess the effect of mutations in this region for their effects on protein stability, ITASSER was used to generate a putative structure of VHL as input for Parameter 6(37).

### Parameters for Algorithmic Assessment

#### Parameter 1: Aggregation Propensity

Aggregation propensity was calculated as previously described(31). A positive aggregation score was was assigned if a given mutation enhanced the hydrophobic character and aggregation propensity of the VHL polypeptide chain. *AGGRESCAN* was used to assess the individual contributions of an amino acid change on the overall hydrophobicity and propensity for aggregation(38).

#### Parameter 2: Protein-Protein Interactions

VHL functions as an E3 ubiquitin ligase when bound to HIF, EloB, EloC, and Cul2(9,10,18,39). To assess the capacity of missense mutation to disrupt these crucial interactions, mutations occurring at positions found to mediate protein interactions with known binding partners were scored positive.

#### Parameter 3: Secondary Structure

VHL consists of three structural domains: an N-terminal random coil region, a beta sheet containing β-domain, and an alpha helical α-domain. Maintaining the secondary structural elements in this region are crucial for pVHL function as pathogenic mutations are less likely to occur in other disordered regions of the protein(28,32). If a missense mutation occurred in a region of secondary structure, it was scored positive for this parameter.

#### Parameter 4: Conformational Flexibility

Flexibility allows a protein molecule to perform its function and bind to substrates and interaction partners(40). The overall flexibility of a given protein is governed by the location of key amino acids within the amino acid sequence. The unique conformational constraint of the proline side chain and the ability to accommodate a *cis-/ trans*-conformation in proteins makes proline a significant contributor to overall protein flexibility and function(41,42). Glycine residues contain a side chain that prevents steric hindrance, increasing the flexibility of a protein(43,44). Finally, cysteine residues are capable of disulfide bonds, which are crucial components of protein stability(45,46). In this analysis, any missense mutation involving changes in proline, glycine, or cysteine residues were scored as positive.

#### Parameter 5: Solvent Accessibility

Replacing surface exposed hydrophilic residues with hydrophobic residues or charged residues with uncharged residues and vice versa can increase the probability of effects on protein-protein interaction and overall protein aggregation(46). In addition, substitution of hydrophobic amino acids for hydrophilic ones within the core of the protein can be thermodynamically unfavorable(47). Finally, the position of charged residues within the protein can be crucial for intramolecular salt bridge formation. Deleterious mutations could destabilize these interactions, thereby destabilizing the protein(48). If a mutation reversed or removed a charge at a given position, replaced a buried hydrophobic residue with a hydrophilic residue, or resulted in a surface exposed hydrophilic residue becoming hydrophobic, it was scored as positive in this parameter.

#### Parameter 6: Protein Stability

Proteins have evolved to fold into specific structures in order to perform their roles in the crowded environment of the cell. We evaluated the effects of missense mutations on the stability of pVHL as destabilizing mutations could prevent proper folding and function. *In vivo* protein folding relies on both thermodynamic and kinetic stability(49,50). The difference in the energy states of the unfolded and that native protein is the thermodynamic stability while kinetic stability refers to the energy barriers that separate any two states of a protein(49,51–54). A missense mutation can alter both the thermodynamic and kinetic stability of a protein indicating a biophysical cause for disease. To determine if overall protein stability was altered via a missense mutation, pVHL missense mutations were assessed using the CUPSAT online prediction server(49). Missense mutations that resulted in a -ΔΔG, i.e., indicating significant changes in overall protein stability, were scored as positive(49).

#### Parameter 7: Post-translational Modifications (PTMs)

Post-translational modifications serve crucial roles on proteins through the covalent addition of small molecules to protein backbones(55). PTMs confer additional specificity to the overall structure and function of a given protein, and contribute to the ability of a protein to interact with different binding partners(55–57). To assess the specific roles of PTMs in our algorithmic assessment of VHL disease, missense mutations that occurred at a position known to be post-translationally modified were positively scored(32).

#### Parameter 8: Translation Rate

A change in the translation rate of a protein can have deleterious effects on folding(58,59). Translation rate is dependent on the codon usage percentage and the number of rare versus common codons in the gene and the subsequent abundance of the corresponding tRNA species. A mutation was scored in this parameter if the mutation change resulted in a translation rate fold change exceeding +2 or −2. Translation rate was calculated using the codon usage tables and tRNA abundances at GtRNAdb(60).

### Overall Score

The overall score given to the multiparametric assessment of each gene mutation was calculated as a sum of the unweighted or weighted scores as described previously(31).

### Parameter Independence and Weighting Strategy

Parameters were tested for independence from one another using Spearmans rho correlation in R. Parameters with rho values < .5 and > −.5 were considered not correlated (Supplemental Table 1). To determine an optimized strategy for weighting score values for each of the parameters, 211 ClinVar mutations were used with their corresponding pathogenicity indicators to develop a pathogenicity index. ClinVar mutations annotated as benign, uncertain significance, or conflicting interpretations were considered “benign” and given a pathogenicity score of 0. Those annotated as likely pathogenic or pathogenic were considered “pathogenic” and given a score of 2. Symphony, an online program to predict the risk of ccRCC in a given VHL mutation, was also used to develop the pathogenicity index by scoring the same 211 ClinVar mutations. Mutations identified as high risk of ccRCC were given a score of 1 while those identified as low risk were given a score of 0. The scores were summed for each mutation, creating a pathogenicity index ranging for 0 to 3 for each of the 211 ClinVar mutations. A chi-square was used to test for dependence of the pathogenicity index score to the unweighted scores of each parameter using R. The resulting p-values were used to set the following cut-offs for our weighting approach. P < .005 was weighted 4. .005 < P < .05 was weighted 3. .05 < P < .5 was weighted 2. Finally, P > .5 was unweighted (i.e. score of 1) (Supplemental Table 2).

**Table 1:**
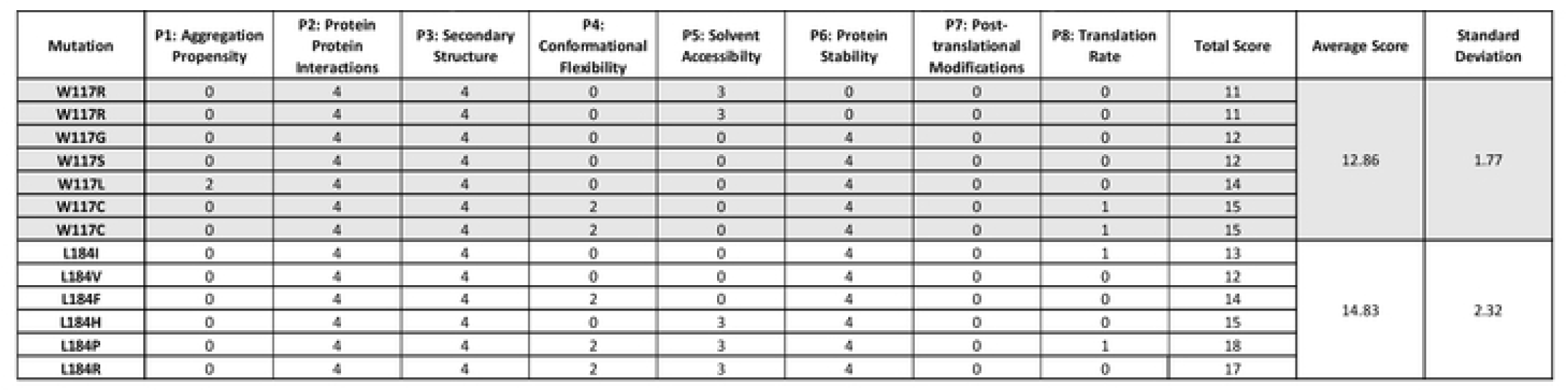
All possible missense mutations at highly destabilizing residues and their corresponding algorithm scores.

**Table 2:**
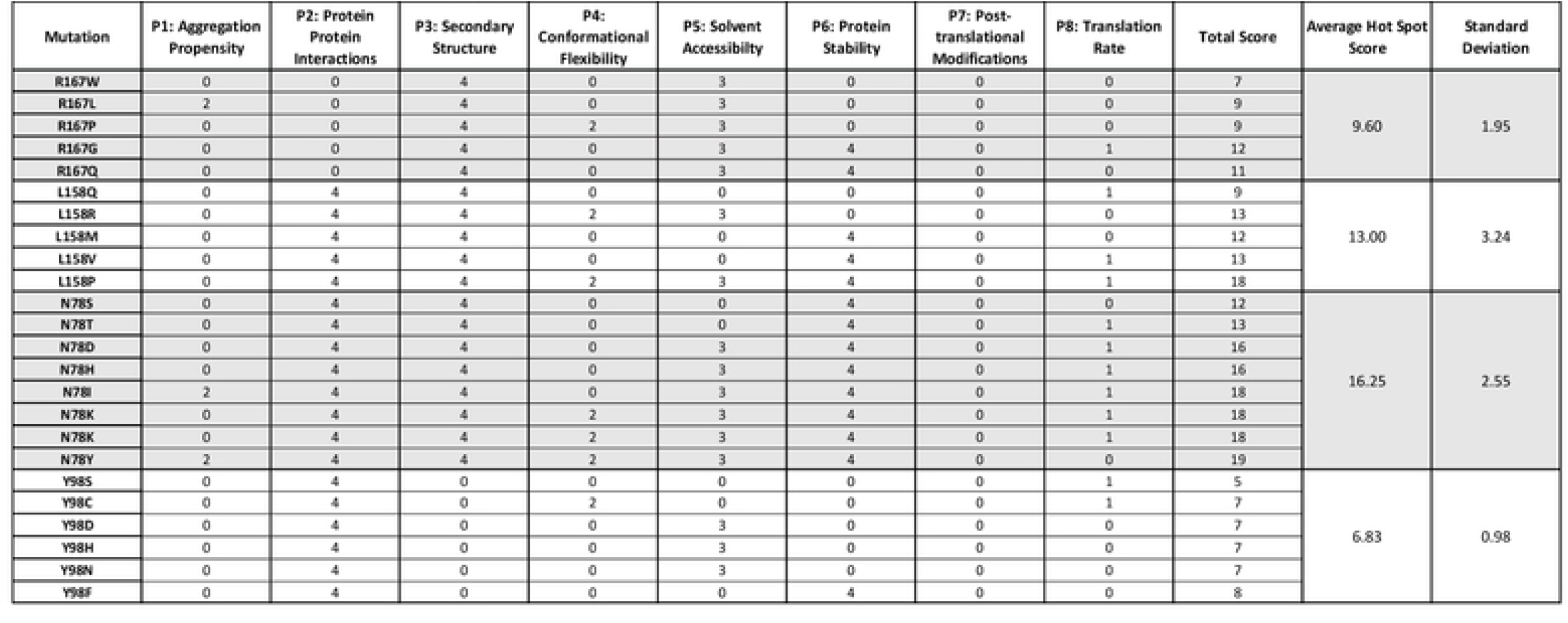
All possible missense mutations at VHL disease associated mutation hot spots and their corresponding algorithm scores.

### Statistical Analysis

All statistical analysis was conducted using GraphPad Prizm (Supplemental File 4). Spearmans rho and Chi-square tests were performed in R (Supplemental Tables 1 & 2).

## Results

Using a set of 285 missense mutations from the ClinVar database and another set of 1380 possible missense mutations (APMM) in pVHL, we began to evaluate the consequences of missense mutations, arising from a SNP. Our multiparametric approach provided a holistic view of the consequences of a mutation on the overall structure and stability of pVHL by evaluating the following parameters: aggregation propensity, protein-protein interactions, secondary structure, conformational flexibility, solvent accessibility, protein stability, post-translational modification, and translational rate.

### Unweighted Scores for all possible mutations and the ClinVar dataset

Using our initial, unweighted approach, in which a scored mutation received a 1 and an unscored mutation received a 0, we obtained a range of values for all missense mutation from 0 to 7 for both the APMM and the ClinVar data sets, indicating no single mutation received a score in all of the 8 parameters (Supplemental Figure 1). Using the ClinVar data set, all of the parameters were determined to be independent of one another (Supplemental Table 1). The average score of the APMM and the ClinVar data sets were 2.7 and 2.8, respectively (Supplemental File 4). Our unweighted approach did not result in significant separations in the benign and pathogenic mutations (means of 2.46 and 3.59, respectively); therefore, we next evaluated the scores using a weighted approach (Supplemental Figure 2A and Supplemental File 4).

**Figure 1:**
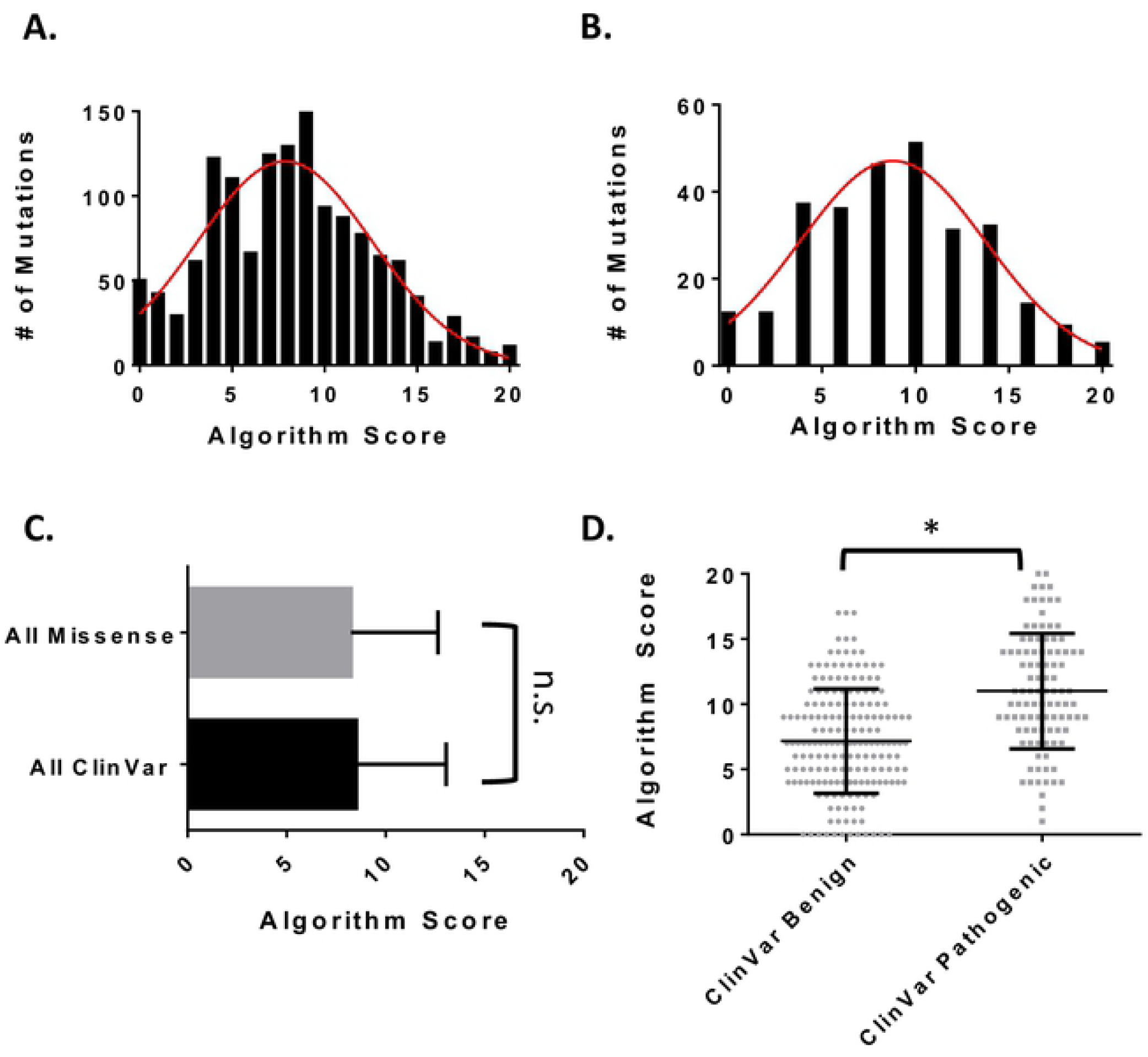
Score distributions for the VHL missense mutations used in the multiparametric approach. **A**. A fitted Gaussian distribution (red) of scores for all 1379 possible missense mutations from a SNP in VHL **B**. A fitted Gaussian distribution (red) of scores for the 285 ClinVar missense mutations used in this study. **C**. Relationship between the All Mutation data set and the ClinVar data set. **D**. Mutation algorithm scores plotted according to their ClinVar pathogenicity. Each dot is a mutation. All error bars represent the standard deviation. A* represents a P < .05 according to a Kolmogorov Smirnov test. All statistics done in Graph Pad Prizm.

**Figure 2:**
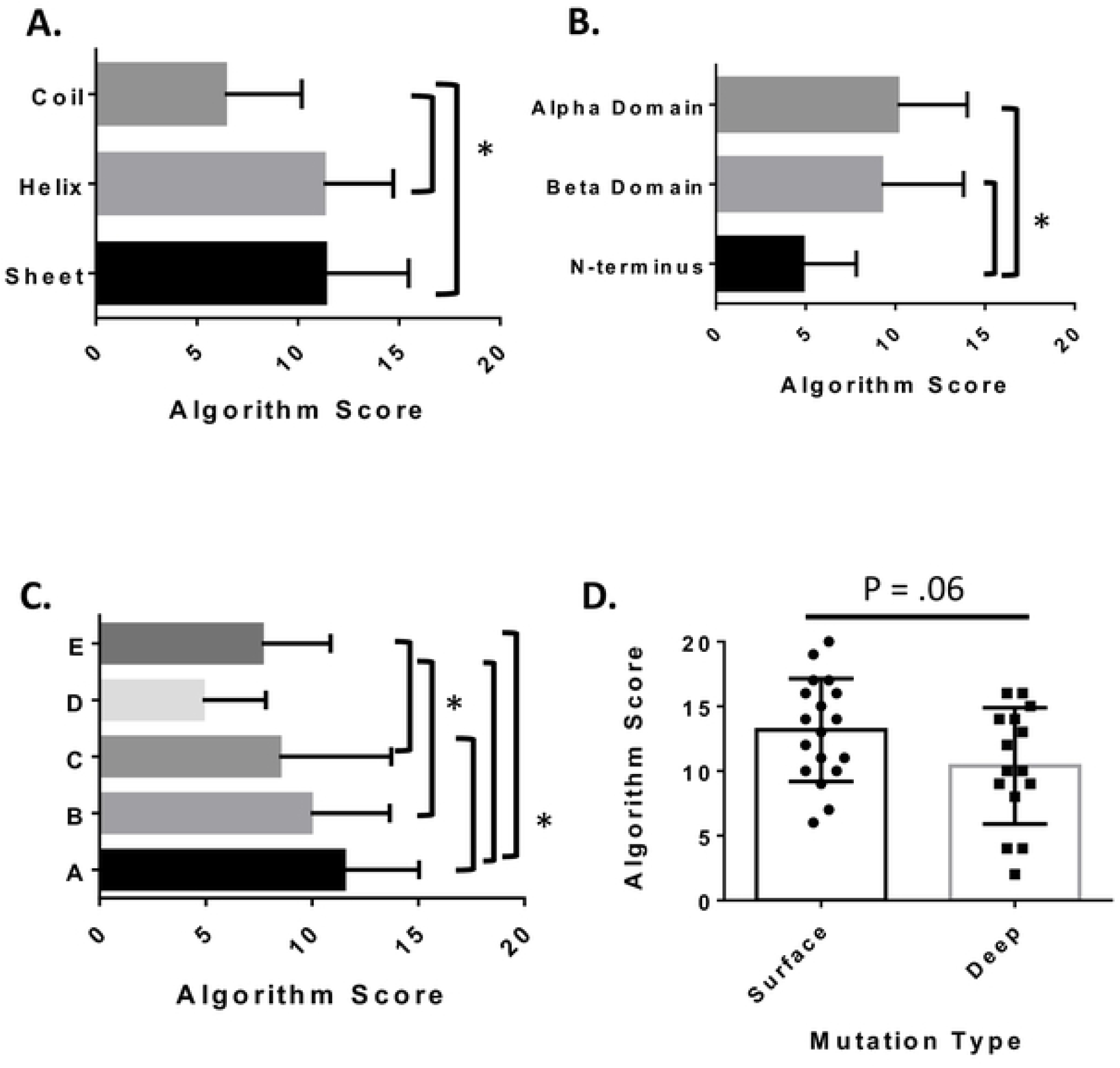
Association of missense mutation algorithm score to its spatial distribution on pVHL. **A**. Algorithm scores for mutations according to secondary structure. **B**. pVHL domain **C**. or pVHL binding interfaces. Significance was determined using an ANOVA or Kruskal-Wallis test and followed up with Tukey HSD or Dunn’s MCT as appropriate. Error bars represent the standard deviation. * represents a significant difference with a p < .05. **D**. Algorithm Score for mutations according to their depth within the structure of VHL. Each dot is a mutation. Error bars represent the standard deviation. * represents a significant difference with a p < .05 as determined by Student’s t-test. All statistics were done using GraphPad Prizm.

### Weighted Scoring Approach

In order to improve our strategy for the algorithmic assessment of missense mutations, a weighting strategy was developed using the pathogenicity indications available on ClinVar and Symphony, an online predictor of ccRCC risk of mutations in VHL. This pathogenicity index was tested for dependence against the unweighted parameters using the chi-square statistic. Weights were then assigned to the parameters according to their resulting p-value (Supplemental Table 2). This new scoring approach resulted in a range of scores from 0 to 20 for both the APMM sand the ClinVar data sets with means of 8.3 and 8.5, respectively (Figure 1 A-B and Supplemental File 4). These populations were not found to be significantly different from one another by a Kolmogorov Smirnov test (P = .91) (Figure 1C). Upon comparing the benign ClinVar mutations to the pathogenic ClinVar mutations, we observed a significant shift in the mean score from 7.2 to 11.0 respectively (Supplemental Figure 2B and Supplemental File 4). This was determined to a be significant difference according to a t-test with a p < .05 (Figure 1D). This ClinVar set was further subdivided into its original ClinVar pathogenicity indications. All of the pathogenic groups (likely pathogenic, likely pathogenic/pathogenic, and pathogenic annotation) showed significant separation from the mutations of uncertain significance (Supplemental Figure 3A). Symphony was also used to determine the risk of ccRCC associated with the ClinVar mutations used in our pathogenicity index. When comparing the risk of ccRCC for the ClinVar mutations, we observed a significant difference in the algorithm score between the mutations identified as high risk (mean score of 10.5) and those identified as low risk (mean score of 7.6) (Supplemental Figure 3B). Our approach to refine the weights of each parameter therefore, was successful in distinguishing populations of pathogenic mutations and benign mutations from those databases listed.

**Figure 3:**
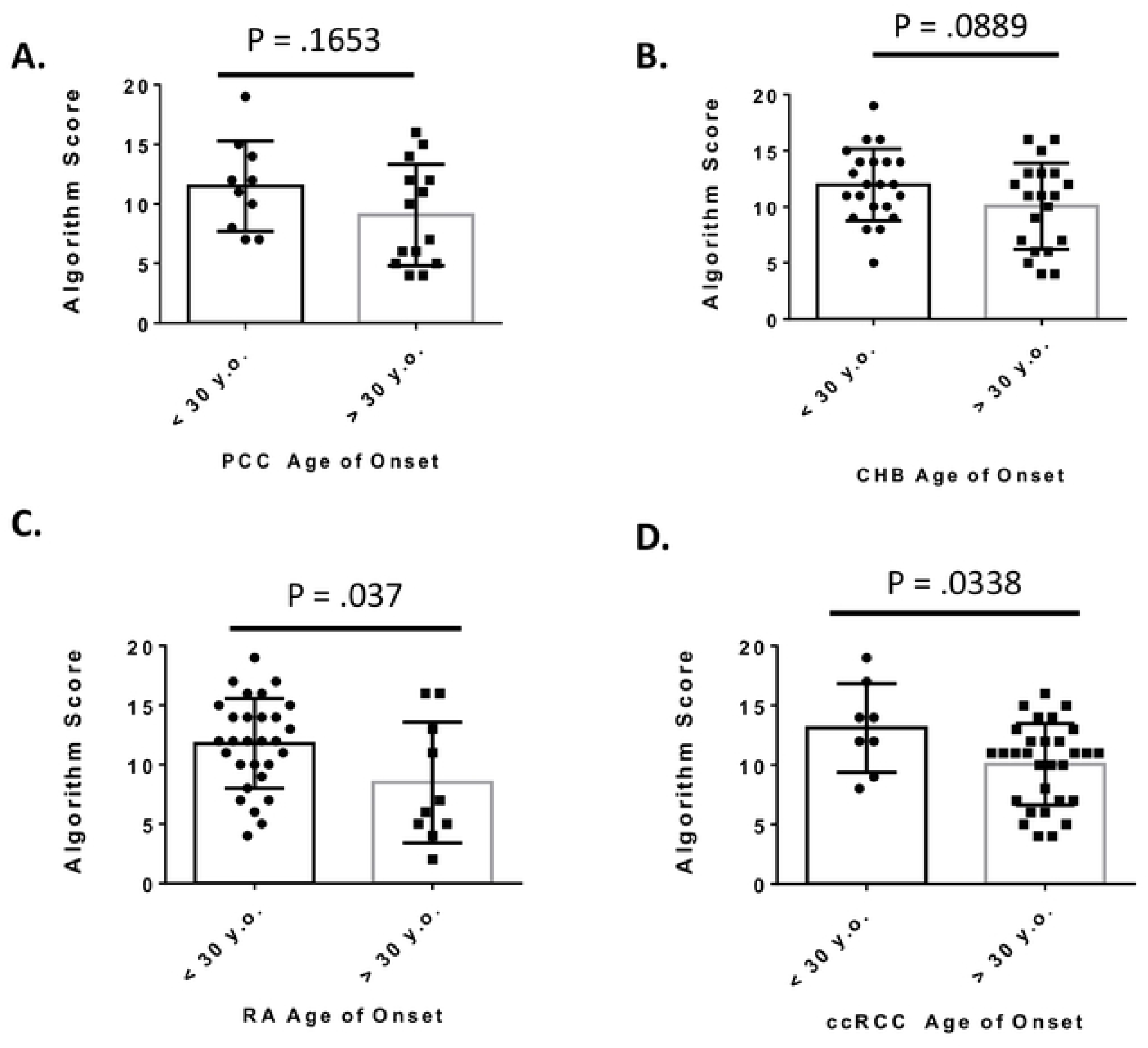
VHL missense mutations algorithm scores associated with onset of the VHL related cancers: **A**. pheochromocytoma (PCC) **B**. central nervous system hemangioblastoma (CHB) **C**. retinal angioma (RA) and **D**. clear cell renal carcinoma (ccRCC). Each dot is the average age of onset for a missense mutation. Error bars represent the standard deviation. P-values were determined using Student’s t-test.

### Algorithm scores according to location within VHL 3-D structure

VHL consists of three structural domains: an N-terminal random coil region, a beta sheet containing β-domain, and an alpha helical α-domain. Pathogenic mutations have been observed to occur at a lower frequency in areas of disorder; therefore, maintaining this arrangement of secondary structure motifs is predicted to be critical for functional pVHL(28,32). We therefore predicted that we should also observe higher algorithm scores in mutations that occur in areas of defined secondary structure. Indeed, mean algorithm scores were significantly higher in regions of helix or sheet compared to random coil regions of the VHL protein (Figure 2A). Overall secondary structure dictates the division of pVHL into three main domains: the α-domain, the β-domain, and the N-terminal coil region(32). We determined that mutations scored higher if they occurred in the α and β domains compared to the N-terminus (Figure 2B).

VHL also consists of five binding interfaces(32,61,62). Interface A is involved in VCB complex formation(9). The HIF-1α binding site is located within interface B(36,39). Cul2 interacts with interface C(32,39). The unstructured N-terminus of VHL is proposed as interface D, though little is known of its binding partners and their importance in the progression of VHL disease(8,32). However, there are residues in interface D that are candidates for phosphorylation by aurora kinase II and casein kinase II(8,32). Finally, interface E, consisting of the helical C-terminus, is predicted to interact with Zinc-finger protein 197 (ZNF-197) and Von-Hippel Lindau Binding Protein 1 (VBP1), a protein chaperone(8,32,63). Due to the importance of each VHL protein interface (A,B,C) in the ubiquitin ligase function of VHL and subsequent HIF regulation by VHL, we expected to observe higher average algorithm scores for mutations occurring within these interfaces (2,39,64,65). When the mean algorithm scores for each of the binding interfaces were compared, we observed significantly higher scores within interface A compared to interfaces C, D, and E (Figure 2C). Interfaces B and C, important binding surfaces for HIF and Cul2 respectively, also had significantly higher algorithm scores than interfaces D and E, which are not involved in VCB complex formation (Figure 2C).

Recent studies into the distribution of mutations within the VHL structure have observed that amino acid changes occurring on the surface of the pVHL are more deleterious for overall function(32,33). These corresponding deleterious genetic mutations are associated with a higher risk of pheochromocytoma (PCC), a cancer of the adrenal glands that causes hormone dysregulation(2,32,33). To determine if these mutations are detected by our algorithmic scoring method, we compared the average algorithm score for mutations at the protein surface and mutations in the protein core (33). Since VHL functions as a scaffold for the assembly of the VCB complex, we would expect that mutations occurring on the surface of the protein, and therefore affecting the binding sites for interaction partners, would result in higher algorithm scores and more severe disease. The p-value (p=.06) indicated that the algorithm scores for comparing the surface versus core-located mutations approached significance at the .05 α-value, suggesting that the observed trend towards a higher algorithm score (mean score = 13.2) in the surface mutations versus mutations occurring deeper in the VHL structure (mean score = 10.4) may have biological importance (Figure 2D). However, additional data and sampling of mutations appropriate for these regional comparisons are needed to improve the statistical score.

### Identification of highly destabilizing and hot spot VHL mutations with algorithmic assessment

We investigated the capacity of our algorithm to identify mutations that have been described as highly destabilizing to VHL(28). Razafinjatovo et al identified W117 and L184 as missense mutation hotspots that can highly destabilize pVHL(28). Our multiparameteric algorithm approach also scored mutations at theses residues considerably higher than the pathogenic mean score of 11 (mean score for APMM at W117 = 12.86 and mean score for APMM at L184 = 14.83), both above the average score for the pathogenic ClinVar mutations (Table 1). Other VHL mutation hot spots, such as L158 and N78 (scores of 13.0 and 16.3, respectively), also scored highly above the average for pathogenic ClinVar mutations; however, R167, another annotated VHL hotspot, received a below average pathogenic score (9.6) (Table 2). Finally, other hotspot mutations, such as Y98 (mean score of 6.8), scored below average for benign scores. Our multiparametric scoring algorithm is designed to provide an evaluative sum of how a given missense mutation will affect the ability of a protein to fold and function properly. In this way, pathogenic mutations such as Y98, with low algorithm scores may not ultimately cause disease phenotypes by destabilizing pVHL protein, but through a more direct local effect that is critical for VHL function and protein interaction. This is likely the case for Y98, located in binding interface B, which is crucial for interaction with HIFα(39,66). Therefore, mutations at these positions (Y98 mutations all score in parameter 2) are sufficient to cause disease through their ability to uniquely affect protein-protein interactions (Table 2). Other studies have found that specific mutations at the Y98 position will cause different VHL cancer phenotypes with Y98H causing type 2B disease and Y98N causing type 2A disease by modulating the efficacy of binding to HIFα(18). Although these kinds of critical mutations (crucial binding site, catalytic abatement, posttranslational substrate) should be taken into account independently from our algorithm, the use of our algorithm scores *combined* with these additional considerations will serve as a valuable comprehensive evaluation for the protein.

### VHL missense mutations score and onset of VHL related cancers

Next, we assessed if our algorithm would be able to identify missense mutations that are more likely to be associated with an early age of onset of VHL-related pathologies. Using published data sets of missense mutations from Chinese patients (available in Peng et al) and another dataset of English patients (available in Ong et al), we compared the algorithm score for 56 missense mutations in early (less than 30 years old) to late (greater than 30 years old) onset of pheochromocytoma (PCC), central nervous system hemangioblastoma (CHB), retinal angioma (RA), and clear cell renal carcinoma (ccRCC)(25,33). For PCC, we observed a shift towards a higher average algorithm score for mutations associated with an early age of onset (11.5) than mutations with a later age of onset (9.1); however, this difference was not statistically significant (p = .1653) (Figure 3A). Similar to PCC, algorithm scores trended towards higher values for early onset of CHB with mean score of 11.9 versus late onset of CHB with a mean score of 10.0; however, this was not a statistically significant difference (p = .0889) (Figure 3B). However, we did see significant differences for the onset of RA and ccRCC (Figure 3C-D). For RA, early onset mutations had an average algorithm score of 11.8 while late onset scores had an average algorithm score of 8.5 (Figure 3C). For ccRCC, early onset mutations had an average algorithm score of 13.1 while late onset mutations had an average algorithm score of 10.1 (Figure 3D).

These data indicate that our algorithm can distinguish more pathogenic mutations from less pathogenic ones that are based on age-related onset of different VHL related cancer types. While not significant at the α-value cut-off at .05, the scores for early age of onset for both PCC and CHB trended towards higher values than the later age of onset. For ccRCC and PA, the scores for early onset versus the scores late onset were significantly higher at α-value cut off of .05. Larger patient datasets from similar studies could be used to further refine our algorithm, and determine significance for both PCC and CHB disease types. Our analysis provide significant support for the use and refinement of *in silico* evaluation of *VHL* mutations and their capacity for large scale protein dysfunction to predict pathogenic outcomes.

## Discussion and Conclusions

Von Hippel-Lindau (VHL) disease is an autosomal dominant hereditary disease that causes a variety of highly vascularized tumors in patients(67,68). While the average life expectancy is around 65 years of age, secondary conditions of tumor development such as blindness or neurological complications can be debilitating. Should these complications go undiagnosed and subsequently untreated, VHL becomes a fatal condition(1,23). Genetic diagnosis of VHL disease provides an early detection method for clinicians to begin surveillance. Computational and biophysical approaches aimed at predicting the severity of a mutation and its deleterious consequences on the function of pVHL can contribute additional information on how the disease might progress. We have provided a multiparameteric algorithmic approach to evaluate the severity of missense mutations in *VHL*. pVHL functions as a scaffold for the creation of the E3 ubiquitin ligase complex for proper regulation of HIF; therefore, our comprehensive evaluation of pVHL misfolding and dysfunction provides a structurally and molecularly informed approach to the prediction of mutation severity.

Our approach was able to distinguish between the populations of benign and pathogenic ClinVar mutations (Figure 1D and Supplemental Figure 2B). We also observed significantly higher algorithm scores for those mutations deemed high risk of ccRCC by Symphony (Supplemental Figure 3B). Taken together, our multiparameteric algorithm can be used to identify pathogenic from benign mutations in pVHL.

pVHL functions as a scaffold for the assembly of the VCB-CR complex(9). Perturbations to its secondary structure and binding capacity can have deleterious effects on the function of this complex, primarily the negative regulation of HIF under normoxia(2,13,16). The N-terminal tail of pVHL is only present in the pVHL30 isoform, with mutations occurring in this region being mostly ranked as clinically benign(32,61). Our algorithm scores also demonstrated significantly lower scores for mutations occurring in the N-terminus compared to the α-or β-domains of the protein (Figure 2B). The N-terminus is predicted to exist as an unstructured/random coil region; therefore, we expected lower average algorithm scores for mutations occurring in the coil regions of pVHL (Figure 2A). Finally, the N-terminus includes binding interface D, one of the five binding interfaces of pVHL, which is not known to interact with proteins crucial for the regulation of HIF (8,32,61). Similar to the N-terminal domain and the random coil regions of pVHL, interface D has the lowest average algorithm score compared to the other binding interfaces (Figure 2C). Interface E consists of the C-terminal helix of pVHL; however, not much is known about its potential binding partners and its involvement in VHL disease(8,32,63). Our data indicate that mutations occurring on interface E are less pathogenic, having a lower algorithm score than mutations occurring on binding surface A or B (Figure 2 C). Binding surfaces A, B, and C are involved in VCB complex formation, HIF, and Cul2 binding, respectively(8,32,61). Mutations that occur in this region are poised to interrupt the protein interactions crucial for HIF regulation, leading to tumor development. This is indicated by their higher average mutations scores of 11.5, 9.9, and 8.5 for surfaces A, B, and C, respectively (Figure 2C and Supplemental File 4). Mutations in binding interface A score significantly higher against all other interfaces, while B and C score significantly higher than the N-terminal interface D (Figure 2C and Supplemental File 4). These observations are consistent with the biological functions of these interfaces in the pathogenicity of VHL disease. Since these surfaces are involved in the formation of the E3 ubiquitin ligase complex, the higher algorithm scores are reflective of the potential dysfunction that result from mutations in these regions. pVHL functions to complex proteins together; therefore, mutations occurring on the surface of the protein, regardless of interface, should be more deleterious to overall function than mutations occurring towards the interior of the protein(33). Using a set of defined surface and deep mutations, our algorithmic approach scored surface mutations higher than deep mutations (Figure 2D). This is in agreement with other studies which found surface mutations to be at a higher risk of developing PCC(33).

VHL is an autosomal dominant hereditary disease putting patients at a lifelong risk of tumor development. Upon spontaneous mutation of the wild-type allele in susceptible tissue types, tumor development begins. A predictive outlook for the onset of VHL related cancers could provide clinicians with a more personalized surveillance strategy when provided with a unique mutation. Using data curated from the literature, our algorithm scored missense mutations associated with an earlier age of onset for RA and ccRCC higher than those associated with a late age of onset (Figure 3C & D)(25,33). While, there was a trend towards higher algorithm scores for the early age of onset of PCC and CHB this was not statistically significant at the α-value cut off of .05 (Figure 3A & B). Our multiparametric method scored W117 and L184, two residues identified as prone to highly destabilizing mutations, with high average scores of 12.84 and 14.83, respectively (Table 1)(28). The approach outlined in this paper can identify mutations that are destabilizing, but this trend was not maintained for all mutations identified as VHL mutation hot spots, such as Y98 (Table 2)(32). Our multiparametric scoring algorithm evaluates the consequences of a missense mutation on the overall stability and folding dynamics of pVHL. Pathogenic mutations with lower algorithm scores, such as Y98, may serve a more direct role in protein-protein interactions or posttranslational modification, may be missed in our algorithm. However, it is interesting to speculate that biochemical studies on clinically identified hotspots that are scored lower in our algorithm, may reveal critical residues for VHL function not previously identified..

Additional clinical data will allow us to iteratively refine our algorithm approach. For example, some same-sense mutations can cause exon skipping in VHL, like the synonymous c.414A>G, p.Pro138Pro mutation(69,70). The dysregulation of splicing creates a truncated protein product consisting of exons 1 and 3. This deleterious variant of pVHL is unable to regulate HIF expression(69,70). For synonymous mutations such as c.414A>G, our algorithmic approach, would give this mutation an overall score of 1, as it can only alter the translation rate of the native codon. In exceptional cases as this clinical mutation, a more detailed understanding of the mechanism of exon skipping could inform future algorithmic approaches for the assessment of exon skipping risk in the *VHL* gene.

We have provided the first comprehensive multiparametric assessment of VHL missense mutations on the function of the VHL protein. Our platform provides the first steps to understand the phenotypic heterogeneity associated with missense mutations in pVHL. We anticipate that our algorithm can undergo iterative refinement as additional clinical data is made available, and the predictive capacity of our approach can be therefore be improved as additional research on VHL is available.

## Author Contributions

Conceived and designed the experiments: FF NS SF SL. Performed the experiments: FF NS SF. Analyzed the data: FF NS SF SL. Contributed reagents/materials/analysis tools: FF SF. Wrote the paper: FF SF SL.

## References

1. Gossage L, Eisen T, Maher ER. VHL, the story of a tumour suppressor gene [Internet]. Vol. 15, Nature Reviews Cancer. 2015 [cited 2019 Sep 11]. p. 55–64. Available from: www.nature.com/reviews/cancer

2. Haase V. The VHL Tumor Suppressor: Master Regulator of HIF. Curr Pharm Des [Internet]. 2009 [cited 2019 Oct 9];15(33):3895–903. Available from: https://www.ncbi.nlm.nih.gov/pmc/articles/PMC3622710/pdf/nihms454216.pdf

3. Latif F, Tory K, Gnarra J, Yao M, Duh FM, Orcutt M Lou, et al. Identification of the von Hippel-Lindau disease tumor suppressor gene. Science (80-). 1993;260(5112):1317–20.

4. Tory K, Brauch H, Linehan M, Barba D, Oldfield E, Katz MF, et al. Specific genetic change in tumors associated with von hippel-lindau disease. J Natl Cancer Inst [Internet]. 1989 Jul 19 [cited 2019 Oct 10];81(14):1097–101. Available from: https://academic.oup.com/jnci/article-lookup/doi/10.1093/jnci/81.14.1097

5. Crossey PA, Foster K, Richards FM, Phipps ME, Latif F, Tory K, et al. Molecular genetic investigations of the mechanism of tumourigenesis in von Hippel-Lindau disease: analysis of allele loss in VHL tumours. Hum Genet [Internet]. 1994 Jan [cited 2019 Oct 10];93(1):53–8. Available from: http://www.ncbi.nlm.nih.gov/pubmed/8270255

6. Iliopoulos O, Ohh M, Kaelin WG. pVHL19 is a biologically active product of the von Hippel-Lindau gene arising from internal translation initiation. Proc Natl Acad Sci U S A [Internet]. 1998 Sep 29 [cited 2019 Oct 10];95(20):11661–6. Available from: http://www.ncbi.nlm.nih.gov/pubmed/9751722

7. Blankenship C, Naglich JG, Whaley JM, Seizinger B, Kley N. Alternate choice of initiation codon produces a biologically active product of the von Hippel Lindau gene with tumor suppressor activity. Oncogene [Internet]. 1999 Feb 8 [cited 2019 Oct 10];18(8):1529–35. Available from: http://www.nature.com/articles/1202473

8. Tabaro F, Minervini G, Sundus F, Quaglia F, Leonardi E, Piovesan D, et al. VHLdb: A database of von Hippel-Lindau protein interactors and mutations OPEN. Nat Publ Gr [Internet]. 2016 [cited 2019 May 21]; Available from: www.nature.com/scientificreports/

9. Cardote TAF, Gadd MS, Ciulli A. Crystal Structure of the Cul2-Rbx1-EloBC-VHL Ubiquitin Ligase Complex. Structure [Internet]. 2017 Jun 6 [cited 2019 Oct 8];25(6):901-911.e3. Available from: https://www.sciencedirect.com/science/article/pii/S0969212617301272?via%3Dihub

10. Ohh M, Takagi Y, Aso T, Stebbins CE, Pavletich NP, Zbar B, et al. Synthetic peptides define critical contacts between elongin C, elongin B, and the von Hippel-Lindau protein. J Clin Invest [Internet]. 1999 Dec [cited 2019 Oct 8];104(11):1583–91. Available from: http://www.ncbi.nlm.nih.gov/pubmed/10587522

11. Stebbins CE, Kaelin WG, Pavletich NP. Structure of the VHL-elonginC-elonginB complex: Implications for VHL tumor suppressor function. Science (80-) [Internet]. 1999 Apr 16 [cited 2019 Oct 8];284(5413):455–61. Available from: http://www.ncbi.nlm.nih.gov/pubmed/10205047

12. Kibel A, Iliopoulos O, Decaprio JA, Jr WGK. Binding of the von Hippel-Lindau Tumor Suppressor Protein to Elongin B and C. Science (80-). 1995;269(September):1444–7.

13. Ohh M, Park CW, Ivan M, Hoffman MA, Kim TY, Huang LE, et al. Ubiquitination of hypoxia-inducible factor requires direct binding to the β-domain of the von Hippel - Lindau protein. Nat Cell Biol [Internet]. 2000 Jul 9 [cited 2019 Oct 10];2(7):423–7. Available from: http://www.nature.com/articles/ncb0700_423

14. Iwai K, Yamanaka K, Kamura T, Minato N, Conaway RC, Conaway JW, et al. Identification of the von Hippel-Lindau tumor-suppressor protein as part of an active E3 ubiquitin ligase complex. Proc Natl Acad Sci U S A [Internet]. 1999 [cited 2019 Oct 10];96(22):12436–41. Available from: www.pnas.org

15. Lisztwan J, Imbert G, Wirbelauer C, Gstaiger M, Krek W. The von Hippel-Lindau tumor suppressor protein is a component of an E3 ubiquitin-protein ligase activity. Genes Dev [Internet]. 1999 Jul 15 [cited 2019 Oct 10];13(14):1822–33. Available from: http://www.ncbi.nlm.nih.gov/pubmed/10421634

16. Maxwell PH, Wlesener MS, Chang GW, Clifford SC, Vaux EC, Cockman ME, et al. The tumour suppressor protein VHL targets hypoxia-inducible factors for oxygen-dependent proteolysis. Nature [Internet]. 1999 May [cited 2019 Oct 11];399(6733):271–5. Available from: http://www.nature.com/articles/20459

17. Hon WC, Wilson MI, Harlos K, Claridge TDW, Schofield CJ, Pugh CW, et al. Structural basis for the recognition of hydroxyproline in HIF-1α by pVHL. Nature [Internet]. 2002 [cited 2019 May 29];417(6892):975–8. Available from: www.nature.com/nature

18. Knauth K, Bex C, Jemth P, Buchberger A. Renal cell carcinoma risk in type 2 von Hippel-Lindau disease correlates with defects in pVHL stability and HIF-1α interactions. Oncogene [Internet]. 2006 Jan 31 [cited 2019 Oct 8];25(3):370–7. Available from: http://www.nature.com/articles/1209062

19. Rechsteiner MP, Von Teichman A, Nowicka A, Sulser T, Schraml P, Moch H. VHL gene mutations and their effects on hypoxia inducible factor HIFα: identification of potential driver and passenger mutations. Cancer Res [Internet]. 2011 [cited 2019 Sep 13];71(16):5500–11. Available from: http://cancerres.aacrjournals.org/

20. Patard JJ, Rioux-Leclercq N, Masson D, Zerrouki S, Jouan F, Collet N, et al. Absence of VHL gene alteration and high VEGF expression are associated with tumour aggressiveness and poor survival of renal-cell carcinoma. Br J Cancer [Internet]. 2009 [cited 2019 Oct 18];101(8):1417–24. Available from: http://www.mlpa.com.

21. Lonser RR, Glenn GM, Walther M, Chew EY, Libutti SK, Linehan WM, et al. von Hippel-Lindau disease. Lancet (London, England) [Internet]. 2003 Jun 14 [cited 2019 Oct 11];361(9374):2059–67. Available from: http://www.ncbi.nlm.nih.gov/pubmed/12814730

22. Binderup MLM. Von Hippel-Lindau disease: Diagnosis and factors influencing disease outcome. Dan Med J [Internet]. 2018 [cited 2019 Oct 11];65(3):5461. Available from: https://ugeskriftet.dk/files/scientific_article_files/2018-08/b5461.pdf

23. Binderup MLM, Jensen AM, Budtz-Jørgensen E, Bisgaard ML. Survival and causes of death in patients with von Hippel-Lindau disease. J Med Genet [Internet]. 2017 Jan 1 [cited 2019 Oct 11];54(1):11–8. Available from: http://www.ncbi.nlm.nih.gov/pubmed/27539272

24. Kai RO, Woodward ER, Killick P, Lim C, Macdonald F, Maher ER. Genotype-phenotype correlations in von Hippel-Lindau disease. Hum Mutat. 2007;28(2):143–9.

25. Peng S, Shepard MJ, Wang J, Li T, Ning X, Cai L, et al. Genotype-phenotype correlations in Chinese von Hippel-Lindau disease patients. Oncotarget. 2017;8(24):38456–65.

26. Gallou C, Chauveau D, Richard S, Joly D, Giraud S, Olschwang S, et al. Genotype-phenotype correlation in von Hippel-Lindau families with renal lesions. Hum Mutat [Internet]. 2004 Sep 1 [cited 2019 Sep 13];24(3):215–24. Available from: http://doi.wiley.com/10.1002/humu.20082

27. Liu SJ, Wang JY, Peng SH, Li T, Ning XH, Hong BA, et al. Genotype and phenotype correlation in von Hippel–Lindau disease based on alteration of the HIF-α binding site in VHL protein. Genet Med [Internet]. 2018 Oct 29 [cited 2019 Sep 13];20(10):1266–73. Available from: http://www.nature.com/articles/gim2017261

28. Razafinjatovo C, Bihr S, Mischo A, Vogl U, Schmidinger M, Moch H, et al. Characterization of VHL missense mutations in sporadic clear cell renal cell carcinoma: Hotspots, affected binding domains, functional impact on pVHL and therapeutic relevance. BMC Cancer [Internet]. 2016 [cited 2019 Jun 17];16(1). Available from: https://bmccancer.biomedcentral.com/track/pdf/10.1186/s12885-016-2688-0

29. Landrum MJ, Lee JM, Benson M, Brown GR, Chao C, Chitipiralla S, et al. ClinVar: Improving access to variant interpretations and supporting evidence. Nucleic Acids Res [Internet]. 2018 Jan 4 [cited 2019 Oct 2];46(D1):D1062–7. Available from: http://academic.oup.com/nar/article/46/D1/D1062/4641904

30. Richards S, Aziz N, Bale S, Bick D, Das S, Gastier-Foster J, et al. Standards and guidelines for the interpretation of sequence variants: A joint consensus recommendation of the American College of Medical Genetics and Genomics and the Association for Molecular Pathology. Genet Med. 2015 May 8;17(5):405–24.

31. Ugrinov KG, Freed SD, Thomas CL, Lee SW. A multiparametric computational algorithm for comprehensive assessment of genetic mutations in mucopolysaccharidosis type IIIA (Sanfilippo Syndrome). PLoS One [Internet]. 2015 [cited 2019 May 20];10(3):121511. Available from: www.hgmd.cf.ac.uk

32. Minervini G, Quaglia F, Tabaro F, Tosatto SCE. Genotype-phenotype relations of the von hippel-lindau tumor suppressor inferred from a large-scale analysis of disease mutations and interactors. PLoS Comput Biol [Internet]. 2019 [cited 2019 May 20];15(4). Available from: https://doi.org/10.1371/journal.pcbi.1006478

33. Kai RO, Woodward ER, Killick P, Lim C, Macdonald F, Maher ER. Genotype-phenotype correlations in von Hippel-Lindau disease. Hum Mutat [Internet]. 2007 Feb [cited 2019 Sep 13];28(2):143–9. Available from: http://doi.wiley.com/10.1002/humu.20385

34. Gallou C, Chauveau D, Richard S, Joly D, Giraud S, Olschwang S, et al. Genotype-phenotype correlation in von Hippel-Lindau families with renal lesions. Hum Mutat. 2004;24(3):215–24.

35. Stebbins CE, Jr WGK, Pavletich NP. Structure of the Complex : Implications for VHL Tumor Suppressor Function. Science (80-). 2009;455(1999):455–62.

36. Min JH, Yang H, Ivan M, Gertler F, Kaelin WG, Pavietich NP. Structure of an HIF-1α-pVHL complex: Hydroxyproline recognition in signaling. Science (80-) [Internet]. 2002 Jun 7 [cited 2019 Oct 8];296(5574):1886–9. Available from: http://www.sciencemag.org/cgi/doi/10.1126/science.1073440

37. Yang J, Zhang Y. Protein Structure and Function Prediction Using I-TASSER. Curr Protoc Bioinforma [Internet]. 2015 [cited 2019 May 29];52:5.8.1-5.8.15. Available from: http://zhanglab.ccmb.med.umich.edu/I-TASSER/download/.

38. Conchillo-Solé O, de Groot NS, Avilés FX, Vendrell J, Daura X, Ventura S. AGGRESCAN: A server for the prediction and evaluation of “hot spots” of aggregation in polypeptides. BMC Bioinformatics [Internet]. 2007 Feb 27 [cited 2019 Oct 3];8(1):65. Available from: http://bmcbioinformatics.biomedcentral.com/articles/10.1186/1471-2105-8-65

39. Hon WC, Wilson MI, Harlos K, Claridge TDW, Schofield CJ, Pugh CW, et al. Structural basis for the recognition of hydroxyproline in HIF-1α by pVHL. Nature [Internet]. 2002 Jun 5 [cited 2019 Oct 8];417(6892):975–8. Available from: http://www.nature.com/articles/nature00767

40. Teilum K, Olsen JG, Kragelund BB. Functional aspects of protein flexibility [Internet]. Vol. 66, Cellular and Molecular Life Sciences. SP Birkhäuser Verlag Basel; 2009 [cited 2019 Oct 3]. p. 2231–47. Available from: http://link.springer.com/10.1007/s00018-009-0014-6

41. Vanhoof G, Goossens F, De Meester I, Hendriks D, Scharpe S. Proline motifs in peptides and their biological processing [Internet]. Vol. 9, FASEB Journal. 1995 [cited 2019 Oct 3]. p. 736–44. Available from: http://www.ncbi.nlm.nih.gov/pubmed/7601338

42. Yaron A, Naider F, Scharpe S. Proline-dependent structural and biological properties of peptides and proteins. Crit Rev Biochem Mol Biol [Internet]. 1993 Jan 26 [cited 2019 Oct 3];28(1):31–81. Available from: http://www.tandfonline.com/doi/full/10.3109/10409239309082572

43. Nick Pace C, Martin Scholtz J. A Helix Propensity Scale Based on Experimental Studies of Peptides and Proteins. Biophys J [Internet]. 1998 Jul [cited 2019 Oct 3];75(1):422–7. Available from: http://www.ncbi.nlm.nih.gov/pubmed/9649402

44. Minor DL, Kim PS. Measurement of the β-sheet-forming propensities of amino acids. Nature [Internet]. 1994 Feb [cited 2019 Oct 3];367(6464):660–3. Available from: http://www.nature.com/articles/367660a0

45. Narayan M. Disulfide bonds: protein folding and subcellular protein trafficking. FEBS J [Internet]. 2012 Jul 1 [cited 2019 Oct 3];279(13):2272–82. Available from: http://doi.wiley.com/10.1111/j.1742-4658.2012.08636.x

46. Pechmann S, Levy ED, Tartaglia GG, Vendruscolo M. Physicochemical principles that regulate the competition between functional and dysfunctional association of proteins. Proc Natl Acad Sci U S A [Internet]. 2009 [cited 2019 Oct 3];106(25):10159–64. Available from: https://www.pnas.org/content/pnas/106/25/10159.full.pdf

47. Dill KA. Dominant Forces in Protein Folding. Biochemistry [Internet]. 1990 Aug 7 [cited 2019 Oct 7];29(31):7133–55. Available from: https://pubs.acs.org/doi/abs/10.1021/bi00483a001

48. Kumar S, Nussinov R. Close-range electrostatic interactions in proteins [Internet]. Vol. 3, ChemBioChem. John Wiley & Sons, Ltd; 2002 [cited 2019 Oct 7]. p. 604–17. Available from: http://doi.wiley.com/10.1002/1439-7633%2820020703%293%3A7%3C604%3A%3AAID-CBIC604%3E3.0.CO%3B2-X

49. Gromiha MM. Prediction of protein stability upon point mutations. In: Biochemical Society Transactions [Internet]. 2007 [cited 2019 May 25]. p. 1569–73. Available from: http://cupsat.uni-koeln.de.

50. Plaza Del Pino IM, Ibarra-Molero B, Sanchez-Ruiz JM. Lower kinetic limit to protein thermal stability: A proposal regarding protein stability in vivo and its relation with misfolding diseases. Proteins Struct Funct Genet. 2000;40(1):58–70.

51. Clark PL. Protein folding in the cell: Reshaping the folding funnel. Trends Biochem Sci. 2004;

52. Baker1 D, Agard DA. Perspectives in Biochemistry Kinetics versus Thermodynamics in Protein Folding [Internet]. Vol. 33, Biochemistry. 1994 [cited 2019 Nov 7]. Available from: https://pubs.acs.org/sharingguidelines

53. Dill KA, Ozkan SB, Scott Shell M, Weikl TR. The Protein Folding Problem.

54. Sanchez-Ruiz JM. Protein kinetic stability. Biophysical Chemistry. 2010.

55. Duan G, Walther D. The Roles of Post-translational Modifications in the Context of Protein Interaction Networks. PLoS Comput Biol [Internet]. 2015 [cited 2019 Nov 8];11(2):1004049. Available from: http://www.uniprot.org/docs/ptmlist

56. Huang JX, Lee G, Cavanaugh KE, Chang JW, Gardel ML, Moellering RE. High throughput discovery of functional protein modifications by Hotspot Thermal Profiling. [cited 2019 Nov 8]; Available from: https://doi.org/10.1038/s41592-019-0499-3

57. Basak S, Lu C, Basak A. Post-Translational Protein Modifications of Rare and Unconventional Types: Implications in Functions and Diseases. Curr Med Chem [Internet]. 2016 Mar 15 [cited 2019 Nov 8];23(7):714–45. Available from: http://www.eurekaselect.com/openurl/content.php?genre=article&issn=0929-8673&volume=23&issue=7&spage=714

58. Tsai CJ, Sauna ZE, Kimchi-Sarfaty C, Ambudkar S V., Gottesman MM, Nussinov R. Synonymous Mutations and Ribosome Stalling Can Lead to Altered Folding Pathways and Distinct Minima [Internet]. Vol. 383, Journal of Molecular Biology. Academic Press; 2008 [cited 2019 Oct 3]. p. 281–91. Available from: https://www.sciencedirect.com/science/article/pii/S0022283608009923?via%3Dihub

59. Clarke IV TF, Clark PL. Increased incidence of rare codon clusters at 5’ and 3’ gene termini: Implications for function. BMC Genomics [Internet]. 2010 Feb 18 [cited 2019 Oct 3];11(1):118. Available from: http://bmcgenomics.biomedcentral.com/articles/10.1186/1471-2164-11-118

60. Chan PP, Lowe TM. GtRNAdb 2.0: An expanded database of transfer RNA genes identified in complete and draft genomes. Nucleic Acids Res [Internet]. 2016 Jan 4 [cited 2019 Oct 7];44(D1):D184–9. Available from: https://academic.oup.com/nar/article-lookup/doi/10.1093/nar/gkv1309

61. Minervini G, Mazzotta GM, Masiero A, Sartori E, Corrà S, Potenza E, et al. Isoform-specific interactions of the von Hippel-Lindau tumor suppressor protein. Sci Rep [Internet]. 2015;5:1–9. Available from: http://dx.doi.org/10.1038/srep12605

62. Minervini G, Panizzoni E, Giollo M, Masiero A, Ferrari C, Tosatto SCE, et al. Design and Analysis of a Petri Net Model of the Von Hippel-Lindau (VHL) Tumor Suppressor Interaction Network. 2014 [cited 2019 May 23]; Available from: www.plosone.org

63. Tsuchiya H, Iseda T, Hino O. Identification of a novel protein (VBP-1) binding to the von Hippel-Lindau (VHL) tumor suppressor gene product. Cancer Res [Internet]. 1996 Jul 1 [cited 2019 Oct 9];56(13):2881–5. Available from: http://www.ncbi.nlm.nih.gov/pubmed/8674032

64. Bader HL, Hsu T. Systemic VHL gene functions and the VHL disease. FEBS Lett [Internet]. 2012;586(11):1562–9. Available from: http://dx.doi.org/10.1016/j.febslet.2012.04.032

65. Metelo AM, Noonan HR, Xiang L, Jin Y, Baker R, Kamentsky L, et al. Pharmacological HIF2α inhibition improves VHL disease-associated phenotypes in zebrafish model. J Clin Invest [Internet]. 2015 [cited 2019 Jun 1];125(5):1987–97. Available from: https://dm5migu4zj3pb.cloudfront.net/manuscripts/73000/73665/cache/73665.3-20150521135435-covered-253bed37ca4c1ab43d105aefdf7b5536.pdf

66. Knauth K, Bex C, Jemth P, Buchberger A. Renal cell carcinoma risk in type 2 von Hippel-Lindau disease correlates with defects in pVHL stability and HIF-1a interactions. Oncogene [Internet]. 2006 [cited 2019 Nov 9];25:370–7. Available from: www.nature.com/onc

67. Maher ER, Iselius L, Yates JRW, Littler M, Benjamin C, Harris R, et al. Von Hippel-Lindau disease: A genetic study. J Med Genet [Internet]. 1991 Jul 1 [cited 2019 Oct 14];28(7):443–7. Available from: http://www.ncbi.nlm.nih.gov/pubmed/1895313

68. Neumann HPH, Wiestler OD. Clustering of features of von Hippel-Lindau syndrome: evidence for a complex genetic locus. Lancet [Internet]. 1991 May 4 [cited 2019 Oct 14];337(8749):1052–4. Available from: https://www.sciencedirect.com/science/article/pii/014067369191705Y?via%3Dihub

69. Flores SK, Cheng Z, Jasper AM, Natori K, Okamoto T, Tanabe A, et al. Synonymous but Not Silent: A Synonymous VHL Variant in Exon 2 Confers Susceptibility to Familial Pheochromocytoma and von Hippel-Lindau Disease. J Clin Endocrinol Metab [Internet]. 2019 [cited 2019 Sep 3];104(9):3826–34. Available from: https://academic.oup.com/jcem/article-abstract/104/9/3826/5426809

70. Lenglet M, Robriquet F, Schwarz K, Camps C, Couturier A, Hoogewijs D, et al. Identification of a new VHL exon and complex splicing alterations in familial erythrocytosis or von Hippel-Lindau disease. Blood. 2018;132(5):469–83.

